# Within-host virus spreading and tolerance to infection are key factors of the *Arabidopsis thaliana* age-dependent susceptibility to viral infections

**DOI:** 10.1101/2023.11.21.568017

**Authors:** Izan Melero, Aurelio Gómez-Cadenas, Rubén González, Santiago F. Elena

## Abstract

*Arabidopsis thaliana* is more susceptible to certain viruses during its later developmental stages. The reasons for this age-dependent susceptibility are not fully understood. Here we explored the possible causes by studying the *A. thaliana* infection response to turnip mosaic virus at three developmental stages: vegetative, bolting and flowering. We found that infected plants at later stages downregulate cell wall fortification genes and that this downregulation facilitates the viral spread and systemic infection. Despite being more susceptible to infection, infected flowering plants are more fertile (*i.e*., produce more viable seeds) than vegetative and bolting infected plants; that is, they have greater fitness than plants infected at these earlier developmental stages. Treatment of postbolting plants with salicylic acid increases resistance to infections at the cost of significantly reducing fertility. Together, these observations suggest a negative trade-off between viral susceptibility and plant fertility. Our findings point towards a development-dependent tolerance to infection.

## INTRODUCTION

Plant susceptibility to infections is influenced by their developmental stage. However, the extent and nature of the age-dependent responses to pathogens vary across different pathosystems (Panter & Jones, 2002; Develey-Rivière & Galiana, 2007). For numerous RNA viruses, mature plants are more susceptible to infection than their younger counterparts (Huang *et al*., 2020; Melero *et al*., 2023). In *Arabidopsis thaliana*, this pattern of susceptibility does not hinge on RNAi-mediated defense responses, suggesting the involvement of other defense mechanisms (Huang *et al*., 2020; DeMell et al., 2023).

Here we sought to better understand how host developmental stages influence the hosts’ responses to viral infection. To do so, we studied the infection of turnip mosaic virus (TuMV; species *Turnip mosaic virus*, genus *Potyvirus*, family *Potyviridae*) in *A. thaliana* HEYNH of the accession Col-0 at three different host developmental stages: (*i*) vegetative juvenile stage, where plants allocate resources to increase their size and mass (prebolting), (*ii*) bolting, an indicator of developmental transition from vegetative to reproductive investment (Pouteau & Albertini, 2009), and (*iii*) flowering, where mature plants allocate resources to reproduction (postbolting). The nine viral lineages studied were derived either from a naïve TuMV isolate (Chen *et al*., 2003) or from an *A. thaliana* preadapted one (González *et al*., 2019) after their evolution in prebolting (vegetative stage), bolting or postbolting (flowering stage) host conditions (Melero *et al*., 2022).

## RESULTS

### Transcriptional and hormonal responses to viral infection are dependent on the host’s developmental stage

To understand how hosts respond to infection at different developmental stages, we conducted a two-step analysis. First, we examined the transcriptional response in prebolting, bolting and postbolting hosts. These hosts were inoculated with the nine viral lineages. A gene ontology (Fig. 1A) analysis of induced and repressed genes indicated that the responses to salicylic acid (SA) were affected; the SA responses were induced on all three developmental stages for the infections with the naïve-derived lineages. We also found that functions related with cell wall biogenesis were repressed in response to infection. This downregulation was observed for all the strains on both bolting and postbolting hosts but was absent in prebolting hosts. Bolting and postbolting hosts shared more similar profiles in terms of the most overrepresented biological categories for the DEGs identified than when comparing with prebolting hosts, which showed clearly distinct functional profiles. However, a principal component analysis (PCA) of the whole transcriptional response showed that the bolting and postbolting plants induced a more similar general response (Fig. 1B). Infection status can be distinguished on the PCA of the transcriptional response, as there is a clear grouping of mock-inoculated plants *vs* infected ones. A clustering of the samples depending on the developmental stage in which they were inoculated was also observed. Particularly, postbolting hosts appeared more distant to the other two developmental stages than when comparing bolting and prebolting juvenile hosts between them.

**Fig. 1.**
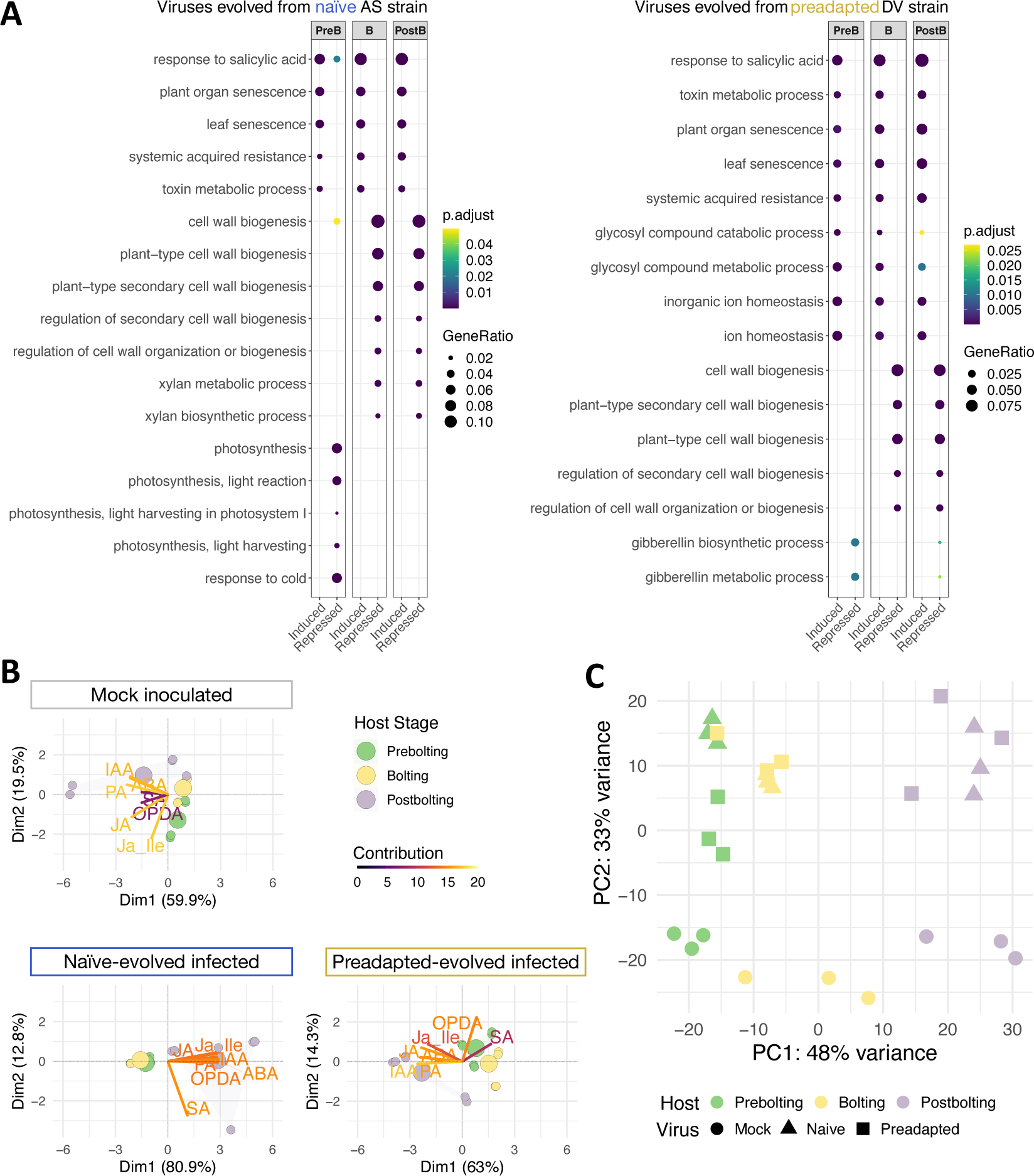
Transcriptional and hormonal response of *A. thaliana* TuMV infection depending on the host developmental stage. **(A)** Gene ontology analysis of differentially expressed genes (DEGs). DEGs upon the infection with the naïve-evolved viruses (left plot) and for the preadapted-evolved ones (right plot). Each column represents the DEGS on each developmental stage - PreB for prebolting, B for bolting, and PostB for postbolting hosts. Induced and repressed genes are showed for each developmental stage. The size of the dots is proportional to the ratio of genes for each biological category, and the color represents the *P* adjusted values. **(B)** Principal component analysis of hormone profiles. Plots are separated for each virus status: mock inoculated plants, naïve-evolved infected plants, and preadapted-evolved infected plants. Dots represent individuals (composed of pools of plants): prebolting plants in green, bolting in yellow, and postbolting in purple. Arrows represent the variables analyzed. The color of the arrow represents the contribution of each variable to the component on the PCA. **(C)** Principal component analysis of the transcriptional response. Samples are differentiated by (*i*) host developmental stage at the time of infection - prebolting in green, bolting in yellow, and postbolting in purple; and (*ii*) inoculum used -circles for mock inoculated plants, triangles for plants infected with the *naïve*-derived viral lineages, and squares for plants infected with the preadapted-derived lineages.

Second, we studied the hormone response to infection. Phytohormones orchestrate plant development and responses to both biotic and abiotic stressors (Zhao & Li, 2021). We measured a comprehensive set of hormones of plants differing in (*i*) their infection status (mock *vs* infected) and (*ii*) the developmental stage in which they were inoculated. We performed a PCA on the data to generate a hormone profile (Fig. 1C). Similar to the PCA of the transcriptional response, the hormonal responses of prebolting and bolting hosts were more alike compared to those of postbolting hosts. Infection led to distinct changes in hormone patterns, notably in SA, which showed a pattern change in plants infected with both viruses. Plants infected with viral lineages evolved from a preadapted ancestor also exhibited altered oxo-phytodienoic acid (OPDA) profiles.

### Faster spread of TuMV in mature plants

As we observed a transcriptional downregulation of cell wall fortification genes in bolting and postbolting plants, we sought to investigate whether the higher viral susceptibility of mature plants could be attributed to this downregulation. We hypothesized that a compromised cell wall might facilitate the virus’ exit from the initially inoculated leaves and its subsequent establishment of a systemic infection. To validate this hypothesis, we studied the time required for the virus to spread within the host by inoculating leaves at the three different developmental stages -prebolting, bolting, and postbolting- and removing the inoculated leaf at various timepoints post-inoculation (Leisner *et al*., 1992; Lafforgue *et al*., 2012). When the inoculated leaves were removed 24- or 48-hours post-inoculation (hpi), we observed that almost no plant at any developmental stage showed signs of infection. However, when we deferred the removal of the inoculated leaves to 72 hpi, there was a noticeable lower incidence of infection in plants at the prebolting (mean frequency ±SE: 0.083 ±0.046) and bolting stages (0.111 ±0.052). The number of infected postbolting plants (0.556 ±0.083) was significantly higher compared to both previous stages (*P* < 0.001 in both cases) (Fig. 2A). This higher rate of infection observed in postbolting plants was slightly lower, yet not significantly different, than observed in postbolting plants that had their inoculated leaves intact (0.625 ±0.099, *P* = 1.000), thereby corroborating the enhanced mobility of TuMV in hosts at this stage. Our results thereby demonstrate that in mature plants TuMV is able to spread faster from the initially inoculated leaf and establish a systemic infection more expeditiously.

**Fig. 2.**
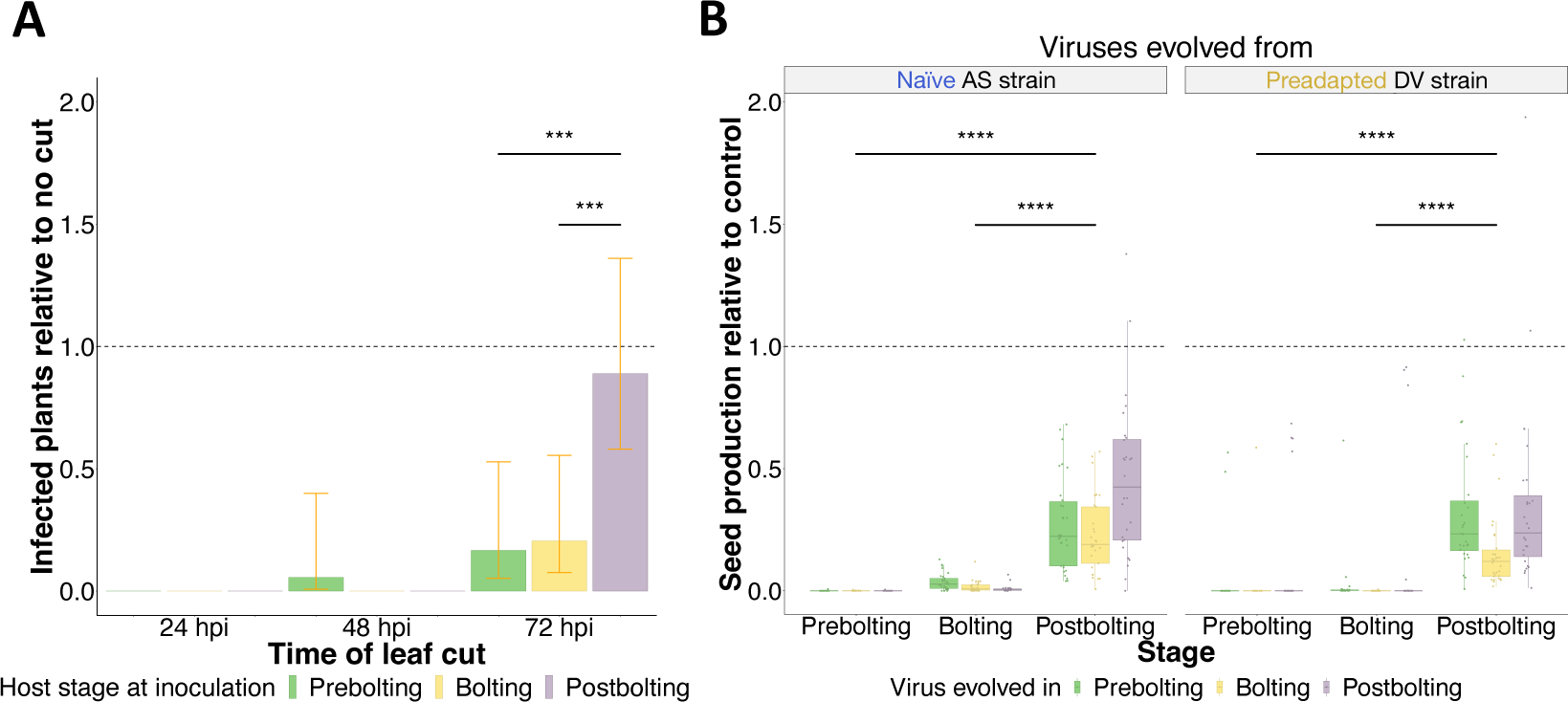
Impact of host developmental stage on virus movement and offspring production. **(A)** Proportion of infected plants after removing the inoculated leaf at different time points (abscissa) in comparation with plants for which the leaf was not removed. Colors represent the host developmental-stage at which the experiment was done (*i*.*e*. at the time of inoculation): prebolting plants in green, bolting plants in yellow, and postbolting plants in purple. Error bars represent ± 95% CI. **(B)** Box plot representation of the number of seeds produced by each infected plant relative to seed production of mock inoculated plants on each developmental stage. Boxes covers the interquartile range and the horizontal lines correspond to the median. On the abscissa, plants are separated by the developmental stage at which they were infected. For each developmental stage, data are grouped depending on the evolved viral inoculum used for the experiment. In green, there are plants infected with viruses evolved on a prebolting stage, in yellow the ones infected with viruses evolved on a bolting stage, and on purple the ones infected with viruses evolved on a postbolting stage. Dots represent individual values of relative seed production for each individual plant. In all cases, asterisks indicate the significance between host developmental stage when inoculated: \*\*\**P* < 0.001; \*\**P* < 0.01; *0.01 < *P* < 0.05.

### Infected mature plants have a higher fitness than plants infected in earlier stages

The plant’s cell wall plays a vital role in seed development. During reproduction, the reshaping of the cell wall becomes crucial for the expansion and steering of the pollen tube, fertilization of the ovule, and the formation of the seed coat (Wolf *et al*., 2012). Mutations that interfere with the biosynthesis or alteration of the cell wall have the potential to hinder these processes and subsequently lower the yield of seeds (Molina *et al*., 2021). Hence, we wondered if infected plants produced different number of seeds depending on the developmental stage at which they were infected. Notice that TuMV is a sterilizing virus (Sánchez *et al*., 2015; Vijayan *et al*., 2017).

We observed clear differences on seed production depending on the developmental stage at which plants were infected (Fig. 2B). Particularly, a very reduced proportion of plants infected at a prebolting juvenile stage produced seeds (9.09%). On the contrary, about half of the plants infected on a bolting stage produced seeds (56.80%) and almost all of the infected plants on a postbolting stage produced seeds (99.42%). For all developmental stages, infected plants produced a smaller number of seeds than their relative mock inoculated plants. Within the infected plants, there was a significant effect of the host stage on the production of seeds: it was higher in postbolting infected plants than in bolting and prebolting hosts, both for viruses evolved from the naïve (*P* < 0.001) or the preadapted (*P* < 0.001) TuMV isolates (Fig. 3B). The developmental stage where viral lineages were evolved in also had an effect on the virus virulence. For lineages derived from the naïve strain, postbolting hosts infected with postbolting-evolved strains had a significative higher production of seeds than when infected with bolting-evolved lineages (*P* < 0.001) or prebolting-evolved ones (*P* < 0.001).

**Fig. 3.**
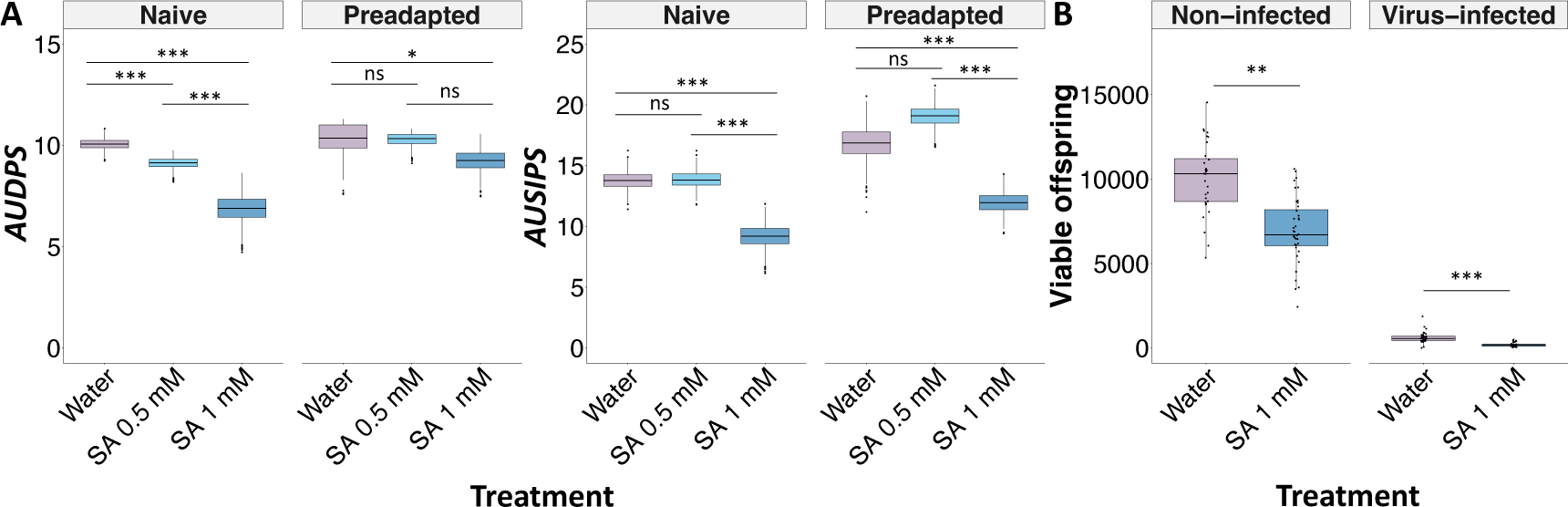
Impact of exogenous application of salicylic acid on susceptibility to infection and production of viable progeny. **(A)** Box plot representation of bootstrapped data of the progression of the infection (*AUDPS*, left panel) or of the symptoms (*AUSIPS*, right panel) of postbolting plants supplemented with water (in purple) or two different concentrations of SA (0.5 mM in light blue, and 1 mM in dark blue). Boxes represent the interquartile range, with the horizontal line corresponding to the median values. Data is divided for plants inoculated with viruses derived from the naïve or the preadapted viral strain. For the naïve virus, the 95% confidence intervals in the *AUDPS* dataset were: for water 10.042 - 10.070, for SA 0.5 mM 9.095 - 9.131, and for SA 1 mM 6.824 - 6.905. For the preadapted virus, they were: for water 10.325 - 10.403, for SA 0.5 mM 10.262 - 10.301, and SA 1 mM 9.202 - 9.267. In the *AUSIPS* dataset, the intervals for the naïve virus were for water 13.716 - 13.806, for SA 0.5 mM 13.815 - 13.900, and for SA 1 mM 9.136 - 9.254. For the preadapted virus, they were for water 16.741 - 16.903,for SA 0.5 mM 18.992 - 19.098, and for SA 1 mM 11.892 - 11.994. **(B)** Number of produced seeds that were viable in non-infected or TuMV naïve-infected plants supplemented with water or 1mM of SA. In all cases, asterisks indicate the significance between host developmental stage when inoculated: \*\*\**P* < 0.001; \*\**P* < 0.01; *0.01 < *P* < 0.05.

Finally, we tested the viability (germination rate) of the seeds produced. Seeds produced by postbolting hosts were more viable: germination of seeds was higher in postbolting hosts (mean ±SD; 0.81 ±0.18) than in bolting (0.57 ±0.05) or prebolting ones (0.30 ±0.01). Hence, the fitness of infected plants is higher if they are infected in later developmental stages.

### Exogenous application of SA increases postbolting plants’ defenses but decreases their fitness

Our study of the transcriptional and hormonal responses suggested an involvement of the SA in the age-dependent susceptibility to TuMV infection. SA plays a crucial role in local and systemic resistance against pathogens, as well as in systemic acquired resistance (SAR), a “whole-plant” response that occurs following an earlier localized exposure to a pathogen (Vlot *et al*., 2009; Gao *et al*., 2015). Hence, we wondered whether by increasing the levels of this hormone, reproductive flowering plants would alter the defense/reproduction balance. To test this, we applied exogenous SA to postbolting plants and inoculated them with both naïve or preadapted TuMV strains (Fig 3A). Disease phenotypes (the progression of infected plants, AUDPS, in the population; or their symptoms, AUSIPS; see methods) indicated that infections progressed slower and were less severe in SA-treated plants in comparison with water-sprayed plants. The mitigation of viral infection in SA-treated plants was more intense in the higher concentration treatment (1 mM).

We then tested the effect of the SA supplementation on the fitness of postbolting plants (Fig 3B). For doing so, we inoculated postbolting plants with the naïve TuMV strain or with buffer (mock), treated with SA 1 mM or water (control), and collected seeds. We found that SA-treated plants produced less seeds than control plants, both in mock (*P* = 0.003) and virus-inoculated plants (*P* < 0.001). These results suggest that the increased susceptibility in postbolting flowering plants could be compensated by a higher production of viable offspring than what it would be produced if the defense response was stronger.

## DISCUSSION

Our results highlight that hosts on later developmental stages suffer from a significant reduction on the expression of genes related with cell wall biosynthesis and metabolism in response to viral infection. The secondary cell wall can play an important role in disease outcomes (Miedes *et al*., 2014). To establish a successful infection, plant viruses not only need to replicate and encapsulate into virions but to spread along the entire host, and for that, they must move from one cell to another through plasmodesmata, and further disseminate to distally located tissues through the phloem (Hipper *et al*., 2013). Plant viruses possess specific proteins, known as movement proteins, which can modulate the size exclusion limit of plasmodesmata (Allie *et al*., 2014) and facilitate the virus’ movement through the plant (Scholthof, 2005; Benitez-Alfonso *et al*. 2010; Niehl & Heinlein, 2011; Schoelz *et al*., 2011). The cell wall identity is a key factor for virus movement (Otulak-Kozieł *et al*., 2018; Koziel *et al*., 2021), as changes on the composition of this barrier can modulate both growth and defense processes (Knox & Benitez-Alfonso, 2014; Xie *et al*., 2018). Some viruses, including TuMV, could move systemically not only through the phloem but also through the xylem (Wan *et al*., 2015; Xue et al., 2023).

Transcriptomic analyses of viral adaptation highlight cell wall modification as an overrepresented category upon viral infection (Hillung *et al*., 2018). We show that cell wall processes are specially repressed on both bolting and flowering plants. We hypothesize that the loosening of cell walls on these hosts may facilitate the deformation of plasmodesmata and enable the faster spread of the virus from one cell to another and, ultimately, to the entire host via the phloem. In line with our results, López-González *et al*. (2021) showed that TuMV-infected plants downregulate genes related with the formation of the secondary cell wall. They found that *IRREGULAR XYLEM 9*, a gene related with xylan synthesis (a key component of the secondary cell wall), was strongly downregulated upon TuMV infection. This downregulation correlated with the reduction of xylan on these infected plants. In our analysis, this gene is among the most downregulated cell-wall related genes in response to all TuMV-strain infections, on both bolting and postbolting plants.

Phytohormones are fundamental players on the interaction between development and immune plant processes (Alazem & Lin, 2015; Zhao & Li, 2021). We observed significant changes in hormone profiles in response to infection across different developmental stages. Interestingly, changes were driven by OPDA and SA. Exogenous application of OPDA and/or SA has been shown to reduce the systemic movement of plant viruses by inducing cell wall fortification (Fernández-Crespo *et al*., 2017). In our work, the exogenous application of SA to postbolting plants decreases their susceptibility to infection as well as their fertility when infected. This result goes in line with our observation of postbolting plants showing higher susceptibility and fertility upon infection, meaning that they showed a tolerance response, understood as a minimized effect of infection on plant fitness (Little *et al*., 2010; Råberg, 2014). Tolerance varies depending on the host and virus genotypes interacting (Montes *et al*., 2020). Here we show that the developmental stage of the host at the time of inoculation is also a key component of its tolerance to viruses. The flowering transition is a critical moment for plant survival, especially for plants that reproduce only once in their lifespan such as *A. thaliana* (Winter *et al*., 2011). Once reproductive development is established, there is a preference of resource allocation at the cost of defense (Winter *et al*., 2011). This can be explained as activating a defense response tends to occur at the cost of reducing growth and reproduction (Bruns, 2016; Karasov *et al*., 2017). The strength and shape of these trade-offs is predicted to drive the evolution towards a lower or higher resistance in younger or older individuals (Buckingham *et al*., 2023). Our observations completely align with these statements: hosts that were infected on their reproductive growth stage produced more viable offspring than those infected on a phase of vegetative growth (which yielded almost no seeds); but in exchange, viruses demonstrated a higher success infecting hosts during the reproductive stage than during the vegetative one. Furthermore, our results evidence that the developmental stage where a virus evolves can drive its virulence: when a naïve strain was evolved in a postbolting stage, the resulting strain was significantly less virulent than the strains evolved in other developmental stages. This reduced virulence of postbolting-evolved strains, as compared to strains evolved in other stages, was not observed for viruses originating from a strain preadapted to prebolting *A. thaliana* plants. This suggests that the evolutionary history of the virus can constrain its evolution towards a lower virulence.

## MATERIAL AND METHODS

### Plant material and growth conditions

*A. thaliana* Col-0 were grown in a climatic chamber under a photoperiod of 16 h light (PAR of 125 µmol m^−2^ s^−1^, produced by a combination of 450 nm blue and 670 nm purple LEDs in a 1:3 ratio) at 24 °C and 8 h dark at 20 °C, 40% relative humidity, in a mixture of 50% DSM WNR1 R73454 substrate (Kekkilä Professional, Vantaa, Finland), 25% grade 3 vermiculite and 25% 3 - 6 mm perlite. Pest management was performed by the introduction of *Stratiolaelaps scimitus* and *Steinernema feltiae* (Koppert Co., Málaga, Spain).

Plants were inoculated at three different developmental stages referred as: prebolting (juvenile plants), bolting (plants on a transition phase between growth stages) and postbolting (mature flowering plants). Under long-day photoperiod conditions, these stages are reached at 18, 25 and 32 days after sowing, respectively. As bolting can be used as indicator of the vegetative to reproductive phase transition (Pouteau & Albertini, 2009), each stage could be associated with different plant stages: vegetative growth, phase transition, and reproductive growth. Following the principal growth stages described by Boyes *et al*. (2001), the inoculated plants correspond to three distinct principal growth stages: prebolting to stage 1.06, bolting to stage 5.10 and postbolting to stage 6.00.

### Virus inoculation

For all experiments and data analysis, except the hypothesis-validation experiments (*i*.*e*., the SA treatment experiment, and the viral spread experiment), we used the TuMV lineages generated in Melero *et al*. (2023). These viruses were evolved on *A. thaliana* hosts on the three aforementioned developmental stages. During the evolution experiment, three lineages were established for each host developmental stage and ten plants were inoculated for each evolved virus. For more details on the evolution experiment, see Melero *et al*. (2023). The original TuMV isolates used for the evolution experiment were: (*i*) TuMV-AS, which came from strain YC5 (GenBank, AF53055.2) originally obtained from calla lily (*Zantedeschia sp*.) and cloned under the 35S promoter and *nos* terminator, resulting in the p35STunos infectious clone (Chen *et al*., 2003) and (*ii*) TuMV-DV, which was obtained from the evolution of the AS isolate for ten passages in 5 weeks-old (prebolting) short-day grown *A. thaliana* Col-0 plants (González *et al*., 2019). For the SA treatment experiment we used the original non-evolved stocks of TuMV-AS and TuMV-DV as viral inoculum. For the viral spread experiment, we only used the original non-evolved stock of TuMV-AS as inoculum.

Inoculations were performed using homogenized virus-infected tissue preserved at −80 °C. The virus inoculum consisted of 100 mg of homogeneous N_2_-frozen infected tissue mixed with 1 mL of phosphate buffer and 10% Carborundum (100 mg/mL). Plants were mechanically inoculated by rubbing 5 μL of the inoculum into three random leaves of the plant. On the viral spread experiment, only one leaf per plant was inoculated.

### Hormone quantification

Aerial tissue was collected and pooled from plants either mock-inoculated or infected at evolution passage five: one pool per each viral lineage (and three pools of mock inoculated plants for each developmental stage). Hormone extraction and analysis were carried out as described by Durgbanshi *et al*. (2005) with few modifications. Briefly, plant tissue was extracted in ultrapure water in a MillMix20 (Domel, Železniki, Slovenia) after spiking with 10 ng of [^2^H_2_]-IAA and 50 ng of the following compounds: [^2^H_6_]-ABA, [^13^C]-SA, [^2^H_3_]-PA, and dihydrojasmonic acid. Following centrifugation, supernatants were recovered, and pH adjusted to 3.0. The water extract was partitioned against diethyl ether and the organic layer recovered and evaporated under vacuum. The residue was resuspended in a 10:90 CH_3_OH:H_2_O solution by gentle sonication. After filtering, the resulting solution was directly injected into an ACQUITY SDS ultra-performance LC system (Waters^TM^, Riga, Latvia). Chromatographic separations were carried out on a reversed-phase C18 column (50×2.1 mm, 1.8-µm particle size; Macherey-Nagel GmbH, Dueren, Germany) using a CH_3_OH:H_2_O (both supplemented with 0.1% acetic acid) gradient. Hormones were quantified using a TQS triple-quadrupole mass spectrometer (Waters^TM^, Riga, Latvia) with two technical replicates performed for each sample. Multivariate analysis was perform using the factoextra package version 1.0.7 (Kassambara & Mundt, 2020) in R version 4.2.0 (R Core Team, 2021) in RStudio version 2022.7.1.554.

### Next-generation sequencing and differential expression analysis

RNA was extracted from evolution passage five (aerial) infected plant tissue using NZY Total RNA Isolation Kit (NZYTech, Lisbon, Portugal). The quality of the RNAs used to prepare RNA-Seq libraries was checked with the Qubit RNA BR Assay Kit (Thermo Fisher, Waltham MA, USA). SMAT libraries, Illumina sequencing (paired end, 150 bp), and quality check of the mRNA-seq libraries were done by Novogene (UK) Co. Ltd. Seventeen bases from the 5’ end and 12 from the 3’ of the reads were trimmed with cutadapt version 2.10 (Martin, 2011). Trimmed sequences were mapped with HiSat2, version 2.1.0 (Zhang *et al*., 2021), to the ENSEMBL release 47 of the Arabidopsis TAIR10 genome assembly. Read counting in features was done with htseq-count, using The Arabidopsis Reference Transcript Dataset (AtRTD2) (Zhang *et al*., 2017) as input annotation file. Differential expression analysis and characterization of differentially expressed genes (DEGs) were done with DESeq2, version 1.24.0 (Love *et al*., 2014), considering only genes having a total of at least 10 reads for each pairwise comparison. For all results, we selected only DEGs of *P* ≤ 0.05 and Ilog_2_-fold changeI > 1.5. Functional profiling was done using the clusterProfiler package version 4.4.4 (Wu *et al*., 2021) with R version 4.2.0 in RStudio version 2022.7.1.554.

### Viral spread experiment

Thirty-six *A. thaliana* plants were virus-inoculated at the three developmental stages studied: prebolting, bolting and postbolting. The inoculated leaf was cut off at either 24, 48 or 72 hpi. As positive controls, 24 plants per developmental stage were virus inoculated without the inoculated leaf being cut off afterwards. As negative controls, 12 plants were mock inoculated, and the inoculated leaf was removed at the same times the virus-inoculated plants’ leaves were cut. Plants were inspected for signs of infection at 14 days post-inoculation (dpi).

### Supplementation with exogenous SA

SA was dissolved in autoclaved Milli-Q water and diluted to achieve concentrations of 0.5 mM and 1 mM. Plants on the three reference developmental stages were inoculated with TuMV-AS and sprayed with (*i*) 0.5 mM SA, (*ii*) 1 mM SA, or (*iii*) water (control plants). Plants were sprayed until they were soaking wet, and substratum was humid. In the experiment done to characterize infection, the exogenous SA was sprayed daily (once a day) from the day plants were inoculated (hours prior to inoculation) until the day prior to their recollection at 14 dpi. In the experiment measuring offspring production, treatments were applied daily for 14 dpi. After this period, treatments were applied once every 3 days for an additional 14 days.

### Infection characterization of SA-treated plants

Upon inoculation, plants were daily inspected for visual symptoms for 14 dpi and phenotyped following a discrete scale of symptoms severity ranging from absence of symptoms (0) to full necrosis of the plant (5) (see Fig. 1 in Butković *et al*., 2021). The infectivity and severity of symptoms data along 14 dpi were used to calculate the area under the disease progress stairs (AUDPS; Simko *et al*., 2012) and intensity progression stairs (AUSIPS; Kone *et al*., 2017) values, respectively, as described in Butković *et al*. (2020). AUDPS and AUSIPS values were computed using the agricolae R package version 1.3-2 (de Mendiburu & Yaseen, 2020) with R version 4.2.0 in RStudio version 2022.7.1.554.

### Seed recollection

Ten plants were inoculated on each developmental stage for each evolved viral lineage. At 14 dpi, the virus-inoculated non-symptomatic plants were discarded, and the symptomatic infected ones were maintained on growing chambers until seed production was reached. Seeds were then recollected for each individual plant separately, and later counted or weighed to obtain an estimation of the number of seeds produced by each plant considering that, on average, a single seed weights ∼20 µg (Qi *et al*., 2017; Sun *et al*., 2017). Mock-inoculated plants were used as a control.

For the plants treated with exogenous SA: 32 plants were mock-inoculated and sprayed with water, 40 were mock-inoculated and sprayed with SA 1 mM, 36 were virus inoculated with TuMV-AS and sprayed with water, and 36 were virus inoculated with TuMV-AS and sprayed with SA 1 mM. The same procedures as previously described were used for seed recollection.

### Germination test

Three blocks of germination tests were done. Seeds from random plants inoculated on the same developmental stage were mixed into a pool, without considering the virus strain used or the stage wherein the virus was previously evolved in. Between 60 and 105 seeds per condition were tested. Seeds were sowed on the same substrate and growing chamber conditions used for plant growing. Viable offspring was counted 10 days after sowing (das).

For the seeds coming from the exogenous SA application treatment: seeds from five random plants were selected from each condition (mock water-sprayed plants; mock SA-treated plants, virus-inoculated water-sprayed plants, and virus-inoculated SA-treated plants). A number between 20 and 50 seeds per plant was evaluated. Seeds were sterilized and sowed on plates containing 2.2 g/L of Murashige-Skoog solid media, 0.5 g/L of ethanesulfonic acid, and 10 g/L of saccharose and maintained in the dark at 25 °C. Viable offspring was counted after 10 das. Wild-type Col-0 seeds were used as control for germination success.

### Statistical analysis

Differences in viral spread rate were tested by fitting the number of infected plants, *I*, to a log-linear factorial model with a logit-link function by means of generalized linear model (GLM). In this case, the model equation reads as logit(*I_ijk_*) ∼ *ɩ* + *D_i_* + *C_j_* + (*D*×*C*)*_ij_* + *ε_ijk_*, where *ɩ* is the grand mean, *D* stands for the developmental stage, *C* for the time at which the inoculated leaf was removed, both factors considered as orthogonal, and *ε_ijk_* represents the Binomial error.

The number of produced seeds (*S*), from plants infected with virus derived from the naïve or the preadapted strain, was fitted to a negative binomial regression with a log-link function using GLM. Host developmental stage where the virus was tested (*T*), host developmental stage where the viral lineage was evolved (*E*), and the ancestral viral strain (*V*) were treated as orthogonal random factors. The full model equation reads as *S_ijkl_* ∼ *σ* + *T_i_* + *E_j_* + *V_k_* + (*T*×*E*)*_ij_* + (*T*×*V*)*_ik_* + (*E*×*V*)*_jk_* + (*T*×*E*×*V*)*_ijk_* +*ε_ijkl_*, where *α* corresponds to the grand mean and *ε_ijkl_* represents the negative binomial error.

For testing the impact of SA treatment on disease phenotypes, the AUDPS and AUSIPS were calculated from the data of a population of 22 inoculated plants per virus strain for the water treatment and 36 inoculated plants per virus strain for each one of the two SA treatments. A bootstrapping method described in Butković *et al*. (2020) was used to estimate 95% confidence intervals. For hypothesis testing, approximated *P* values in pairwise comparisons were obtained by calculating the proportion of overlap of the 95% confidence intervals. *P* values were further adjusted using the sequential Bonferroni’s correction to account for multiple comparisons.

Finally, the number of viable progeny (*R*), was fitted to a GLM with a Gamma distribution and a log-link function in which the treatment (*M*) and the infection status (*F*) were treated as orthogonal random factors. The full model equation reads as *R_ijk_* ∼ *ρ*+ *M_i_* + *F_j_* + (*M*×*F*)*_ij_* +*ε_ijk_*, where *ρ* corresponds to the grand mean and ε*_ijk_* represents gamma-distributed errors.

In all analyses, *post hoc* pairwise comparisons were done using the sequential Bonferroni’s method. Computations were done using SPSS version 28.0.1.0 (IBM, Armonk NY, USA) or R version 3.6.1 within the RStudio version 1.3.1093.

## ACKNOWLEDGMENTS

We thank Francisca de la Iglesia and Paula Agudo for excellent technical assistance. We also thank María J. Olmo-Uceda for great advice and guidance on RNA-seq analysis. This research was supported by grants PRE2020-094661 funded by MCIN/AEI/10.13039/501100011033 and by “ESF investing in your future” (I.M.), PID2022-136912NB-I00 funded by MCIN/AEI/10.13039/501100011033 and by European Union NextGeneration EU/PRTR and CIPROM/2022/59 funded by Generalitat Valenciana (S.F.E), and by an EMBO Postdoctoral Fellowship (ALTF 311-2021) (R.G.).

## AUTHORS’ CONTRIBUTIONS

I.M., data curation, formal analysis, investigation, visualization, writing-original draft. A.G-C., formal analysis, investigation. R.G., conceptualization, formal analysis, investigation, visualization, supervision, writing-original draft, writing-review and editing. S.F.E., conceptualization, formal analysis, funding acquisition, project administration, resources, supervision, writing-review, and editing. All authors gave final approval for publication and agreed to be held accountable for the work performed therein.

## COMPETING INTERESTS

We have no competing interest.

## Notes

### Competing Interest Statement

The authors have declared no competing interest.

